# A Novel Compound, ML336, Inhibits VEEV Replication by Interfering with Viral RNA Synthesis

**DOI:** 10.1101/343228

**Authors:** Andrew M. Skidmore, Robert S. Adcock, Jasper Lee, Colleen B. Jonsson, Jennifer E. Golden, Dong-Hoon Chung

## Abstract

Venezuelan equine encephalitis virus (VEEV) is an alphavirus that is endemic to Central and South America. VEEV is known to cause periodic outbreaks of encephalitis in both humans and equids. There are currently no treatments or preventatives for VEEV disease. Our group has previously reported on the development of a novel VEEV inhibitor, ML336, which showed a potent antiviral effect in cell culture models. However, the mechanism of action had yet to be elucidated. Based on the discovery of mutations conferring resistance within nonstructural proteins, we hypothesized that ML336 inhibits viral RNA synthesis. We found that ML336 was able to inhibit VEEV RNA synthesis with an IC_50_ value of 1.1 nM in a metabolic labelling assay. ML336 marginally affected cellular transcription at levels 20,000-fold above the IC_50_, and did not show any cytotoxicity up to 50 µM. Using a combination of fluorography, strand-specific qRT-PCR, and a metabolic labelling assay, we found that ML336 inhibits the synthesis of all forms of VEEV RNA. Structural analogues of ML336 showed a correlation between their RNA synthesis inhibitory activity and their antiviral activity in cells, leading us to propose that the primary mechanism of action of this class of compounds is viral RNA synthesis inhibition. The activities of ML336 were highly specific to VEEV, without measurable activity against Chikungunya virus. ML336 was efficacious even in a cell-free viral RNA synthesis assay, suggesting a direct interaction with viral proteins.

**Importance:** Venezuelan equine encephalitis virus (VEEV) is a pathogenic alphavirus that circulates in the Americas which can cause a lethal encephalitis in humans and equids. There are currently no licensed treatments or vaccines for VEEV. Due to the high potential for aerosol infection and severe outcomes, it is classified as an NIAID Category B agent. To address the unmet need for VEEV antivirals, we continue to advance a novel amidine compound, ML336, through medicinal chemistry and mechanism of action (MOA) studies. Here, we present the molecular MOA by which ML336 inhibits VEEV replication using cellular and biochemical approaches. Our data suggest that ML336 is a direct-acting antiviral that inhibits viral RNA synthesis by interfering with the viral replicase complex. Our studies provide new insights into approaches for the development of novel RNA virus replication inhibitors and the molecular mechanism of alphavirus RNA synthesis.

## Introduction

Venezuelan equine encephalitis virus (VEEV) is a single-stranded, positive-sense, RNA virus belonging to the family *Togaviridae*, which includes other medically important mosquito-borne viruses such as Chikungunya virus (CHIKV), and Eastern and Western equine encephalitis viruses (EEEV and WEEV, respectively)(1). VEEV, WEEV, and EEEV are closely related, sharing a recent common ancestor as determined by phylogenetic analysis (2), and can cause encephalitis in humans and equids. For VEEV, approximately 1% of patients suffer from encephalitis, leading to death in about 10% of encephalitic patients (1).

These encephalitic alphaviruses have caused periodic outbreaks in the Americas throughout the 20^th^ century, and of these viruses, VEEV has the most significant public health burden. Historically, large VEEV outbreaks occur about every 15 - 20 years, typically affecting thousands of equids and hundreds of humans. For example, between 1962 and 1972 in Central America (3), over 109,000 human cases of VEEV were reported, with nearly 1,000 neurological cases and over 500 associated human fatalities. These outbreaks also caused a significant burden to agriculture with over 800,000 reported deaths of VEEV infected equids.

In addition to large natural outbreaks, VEEV poses additional threats to the public. Historically VEEV has been developed as an agent of biological warfare and still has potential as a biological weapon. Therefore, VEEV is classified as a Select Agent by the Centers for Disease Control and Prevention and United State Department of Agriculture (4). There are currently no licensed treatments or vaccines for VEEV or any other alphavirus infections in humans. Due to the stochastic nature of VEEV outbreaks, therapeutics could be crucial for disease control.

There have been several attempts to develop treatments for VEEV infection using cell culture and animal models with current FDA-approved antivirals; however, those efforts resulted in only limited success (5, 6). Other discovery efforts using nucleoside analogues or non-nucleoside analogues showed only moderate effects against VEEV (7, 8). Currently, only supportive care is available for those suffering from VEEV infection.

To address the unmet need for VEEV therapeutics, we developed and employed a high-throughput screen that led to the discovery of a novel quinazolinone hit compound (CID:15997213) (9). This compound showed significant promise as a small molecule inhibitor of VEEV infection with an EC_50_ of 1.9 µM in a cell culture assay and 100% protection at a dose 50 mg/kg in a lethal VEEV mouse model (10). Refinement of this structure led us to develop a novel amidine compound, ML336, using medicinal chemistry approaches (Fig 1 A) (11). ML336 showed a potent and specific, anti-VEEV activity *in vitro* with an EC_50_ of 32 nM, and titer reduction greater than 7.2 logs at 5 µM (11, 12). ML336 has also been shown to protect mice from a lethal VEEV challenge without any toxicity at the examined doses (11).

**Figure 1.**
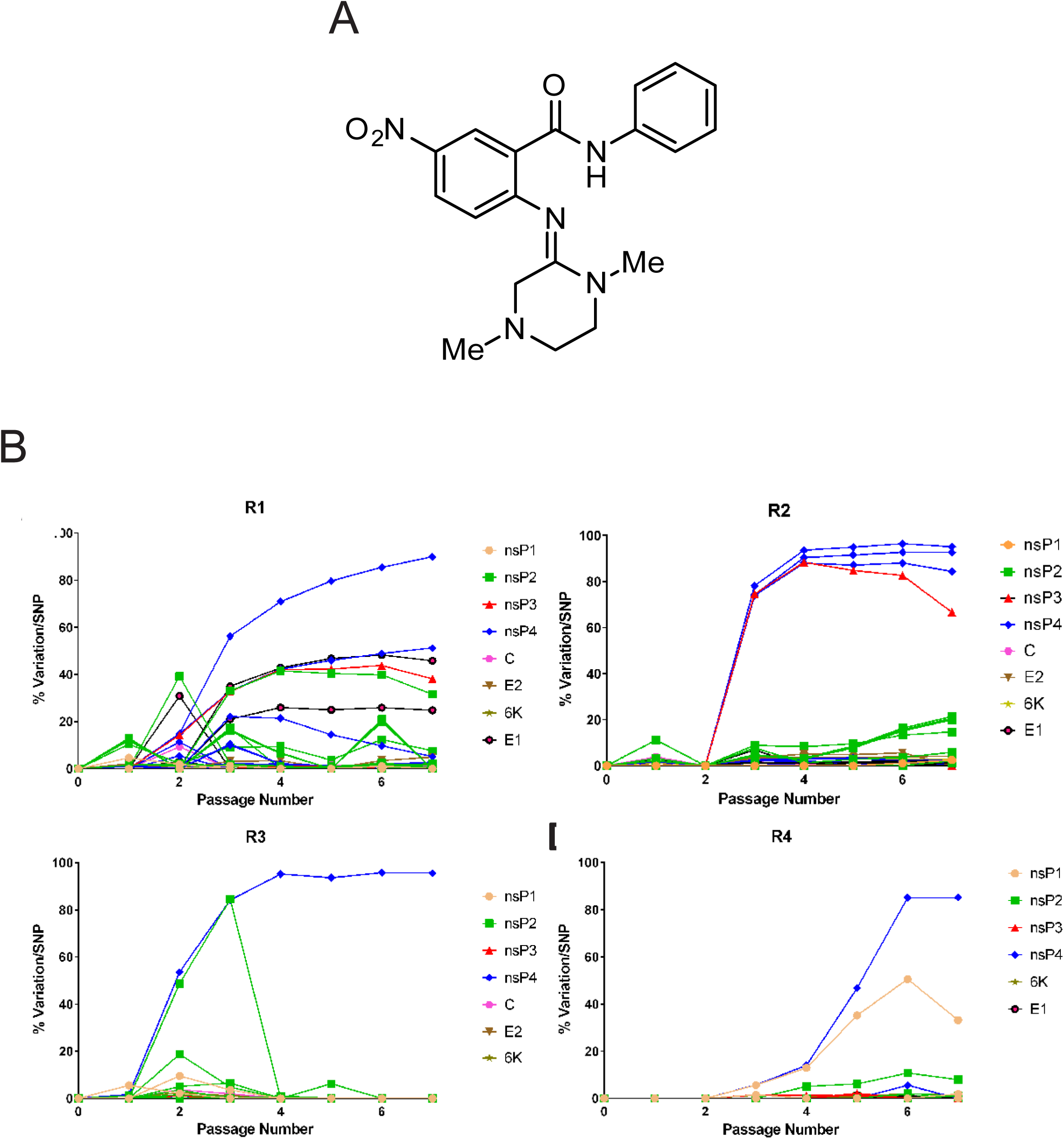
Percentage of variations per SNP in population per passage with ML336. A) The chemical structure of ML336, previously published (11). B) Each replicate (R1, R2, R3, and R4) was passaged separately seven times through Vero76 cells with increasing concentrations of ML336. For each successive passage, the ML336 concentration was doubled, starting at 50 nM, and ending at 3200 nM on passage 7. In each graph, each mutation is individually plotted, comparing each SNPs percent frequency at the passage number in which it appeared. Mutations are color-coded by gene.

While ML336 effectively inhibits VEEV replication, as determined by titer reduction assay, the precise mechanism of action has not been defined. Our previous studies have indicated that both the hit quinazolinone compound (Pubchem CID:15997213) and ML336 might inhibit VEEV replication through non-structural proteins 2 and 4 (nsP2 and nsP4) in the middle phase of replication (10). Resistant mutant selection studies using the quinazolinone compound mapped resistant mutations within two regions; 1) an area upstream to all known activity domains of nsP2 (Y102C, D116N, E117V, and E118V in nsP2) and 2) an area upstream to the RNA-dependent RNA polymerase domain of nsP4 (Q210K and Q210R in nsP4) (10, 13-18). Currently the biological functions of these regions have not been characterized (14).

As the primary function of alphavirus nsPs is the replication of the viral genome, we hypothesized that ML336 inhibits viral replication by interfering with viral RNA synthesis (1, 19, 20). After the initial translation of the incoming viral RNA, the non-structural polyprotein undergoes cleavages mediated by protease activity of nsP2 (18, 21). Intermediate complex nsP123/4 produces negative sense template RNA (22, 23). The mature replicase complex, nsP1/2/3/4, synthesizes the positive sense viral genomic and 26S subgenomic RNAs (23, 24). This viral RNA synthesis occurs in spherules, micro-invaginations on intracellular organelles, in the infected cells (25, 26).

Here, we demonstrate that ML336 efficiently inhibits VEEV replication by interfering with viral RNA synthesis. ML336 prevented the production of VEEV RNA in infected cells as well as in an *in vitro* RNA synthesis assay, indicating that ML336 is a direct-acting antiviral against VEEV. We show that ML336 is a promising small molecule lead compound for VEEV therapeutic development and as a molecular probe to study alphaviral replication.

## Materials and Methods

### Cell culture and viral strains

Baby hamster kidney (BHK) clone 21 cells (ATCC CCL-10) and Vero 76 (ATCC^®^ CRL-1587™) were maintained in Modified Eagle’s Medium with Earle’s Balanced Salt Solution and L-glutamine (MEM-E) supplemented with 10% fetal bovine serum (FBS) (Corning CellGro). Cells were maintained at 37 °C in humidified incubators with 5% CO_2._ VEEV strain TC-83 (gift of Dr. Connie Schmaljohn, USAMRIID) was used for this study. Infections were carried out using a virus infection medium (Modified Eagle’s Medium with Earle’s Balanced Salt Solution supplemented with 1x GlutaMAX, 25 mM HEPES, and 10% FBS. For Chikungunya virus (CHIKV) experiments, CHIKV strain 181/25 (BEI Resources NR-13222) was used.

### Isolation of ML336 resistant viral strains

Vero76 cells were cultured in 6-well plates in Dulbecco’s modified Eagle’s medium (DMEM) supplemented with 10% FBS and 5 mM penicillin/streptomycin at 37°C, 5% CO_2_. Cells were initially infected with TC-83 at an MOI of 1 and 50 nM ML336 was added to cells following infection for 1 h (passage one). Supernatant was collected at 24 hours post-infection and 250 μL was taken for the subsequent passage. The concentration of ML336 at was doubled for each passage (i.e., 50, 100, 200, 400, 800, 1600, 3200 nM) for a total of seven passages

Total RNA was isolated from viral supernatants using Trizol LS. cDNA was synthesized from purified RNA using random hexamer primers and the SuperScript IV First-Strand Synthesis System (Thermofisher). cDNA was amplified through 30 rounds of PCR, using the Phusion High-Fidelity Kit (Thermofisher F553L) and a set of 25 primer pairs for two times coverage of the whole genome, generating PCR fragments either 500 or 1000 base pairs in length. PCR products were purified using the Wizard SV Gel and PCR Clean-Up System (Promega A9282). Libraries for next-generation sequencing were prepared using the Nextera XT kit (Illumina), then sequenced on either the Illumina MiSeq or NextSeq 500. Data was analyzed using CLC Genomics Workbench v.10 (Qiagen). Reads were filtered with a Phred score < 20, mapped reads to the seed stock consensus sequence, and identified variants that appeared above 1% in all reads at positions with a coverage of at least 400.

### Western blot analysis

Cells on culture plates were lysed using a cell lysis buffer containing sodium dodecyl sulfate (SDS) and 42 µM of dithiothretol (DTT) and then DNA was sheared using a needle and syringe, with additional sonication as necessary. After denaturation at 100 °C for ten minutes, sample was loaded onto a gradient gel (4-20%, GenScripts) and run at 140 V for 90 minutes in 1X MOPS-SDS buffer (GenScripts). Proteins were transferred to a PVDF membrane. The membrane was blocked and probed using antibodies for the indicated proteins. Anti-nsP2 (Clone no. 8A4B3) was generated as a custom mouse monoclonal antibody from GenScripts using bacterially expressed recombinant nsP2 protein. Beta-actin was detected using an antibody directly linked to horse radish peroxidase (HRP) (Cell Signaling). Anti-nsP2 was detected using (HRP)-conjugated rabbit anti-mouse IgG antibody. Images were developed using ECL reagent and captured using an Azure Biosystems c300 imaging system.

### Analysis of viral RNA synthesis *in vivo* by metabolic labelling with ^3^H-uridine

Cells were infected with virus at a multiplicity of infection (MOI) of ten on ice for one hour. Then the cells were washed with ice-cold 1X phosphate buffered saline (PBS) and then transferred to a 37 °C CO_2_ incubator to initiate the replication (T=0). At 6 hours post-infection (HPI), cells were washed and pulsed with virus infection media containing actinomycin D (act D) (1 µg/mL, Sigma Aldrich), tritium-labelled uridine (3HU) (5 µCi/mL, Perkin Elmer), and ML336 at various concentration for two hours. Total RNA was isolated from the cells using RNAzol RT according to manufacturer’s instructions (Molecular Research Center). Total RNA was mixed with 10 mL liquid scintillation cocktail (BETA BLEND, MP Biochemicals) and the radioactivity was measured using a Perking Elmer Tri-Carb 2910 TR liquid scintillation counter. For CHIKV, infection proceeded to 8 HPI for 3HU pulse labelling. For compound treatment, ML336 dissolved in DMSO was added in the 3HU labelling mixture with a final DMSO concentration of 0.25%. For cyclohexamide (CHX) treatment, cells were treated with 3HU labelling mixture containing CHX with a final concentration of 8.8 µg/mL. Fluorography of 3HU-labelled viral RNA was performed following a protocol from the John Aris lab (27).

### Strand-specific quantitative real-time PCR (qRT-PCR) of VEEV RNA

Detection of positive and negative sense, genomic viral RNA was carried out using a strand-specific qRT-PCR method adapted from a published paper (28). Briefly, cDNA was generated using tagged primers; **GGC AGT ATC GTG AAT TCG ATG C***GGCGACTCTAACTCCCTTATTG* and **GGCAGT ATCGTGAATTCGATGC***CTGACCTGGAAACTGAGACTATG* for detecting positive-sense and negative-sense RNA, respectively. The generated cDNA was then used in qRT-PCR using TaqMan chemistry with a strand-specific primer set. For positive-sense RNA detection, forward primer 5’aataaatcataa *CTG ACC TGG AAA CTG AGA CTA TG* 3’ and reverse primer 5’aataaatcataa **GGC AGT ATC GTG AAT TCG ATG C** 3’ were used. For negative-sense RNA detection, forward primer 5’aataaatcataa *GGC GAC TCT AAC TCC CTT ATT G* 3’, and reverse primer 5’aataaatcataa **GGC AGT ATC GTG AAT TCG ATG C** 3’ were used. A fluorescent probe (/56-FAM/TCC GTC AAC /ZEN/CGC GTA TAC ATC CTG /3IABkFQ) was used for both analyses. Lowercase sequences are additional sequence added to increase primer identification, *ITALIC* sequences are those that are specific for viral RNA, **BOLD** sequences are those that are used to identify only those cDNA sequences that were produced due to primer binding.

### Enrichment of viral replicase complexes from infected cells

VEEV replicase complexes were isolated according to the protocol published by Barton et al (29). Cells were infected with VEEV TC-83 at 10 MOI and incubated for 6 hours. Then, cells were washed with ice-cold, sterile PBS and the cells were incubated in hypotonic RS buffer (10 mM NaCl, 10 mM Tris-HCl, pH= 7.8) supplemented with protease inhibitor cocktail (Research Products International Protease inhibitor cocktail III) on ice for 15 minutes. Cells were scraped into buffer and thoroughly homogenized using a Dounce homogenizer. The nuclei were removed by centrifugation at 900 x g for 10 minutes at 4 °C. Supernatant containing the cytoplasmic fraction was transferred to microcentrifuge tubes and centrifuged at 15,000 x g for 20 minutes at 4 °C. The supernatant (S15 fraction) was removed and pellets (P15 fraction) were suspended in RS buffer supplemented to 15% glycerol for storage at −80 °C.

### *In vitro* viral RNA synthesis assay

VEEV viral RNA synthesis assay was adapted from Barton et al (29). Ten microliter of P15 fraction enriched for VEEV viral replicase complexes, which is equivalent of approximately 1.25 × 10^6^ infected cells, was combined with a same volume of a RNA synthesis mix (100 mM Tris-HCl pH 7.8, 100 mM KCl, 20 µg/mL act D, 20 mM DTT, 10 mM creatine phosphate, 50 µg/reaction creatine phosphokinase, 4 mM of ATP, GTP, and UTP, 20 µM CTP, 12 mM MgCl2) on ice and 1 µL of SUPERase In RNase inhibitor (Ambion), 5 µg of yeast tRNA (Ambion), and 5 µCi of [α-^33^P]-CTP (Perkin-Elmer) were added per reaction. After an incubation at 37 °C for 90 minutes, RNA was isolated from each reaction using RNAzol RT and RNA mini prep kit columns (Zymo Research) according to manufacturer’s instructions, with an additional wash step before elution. For compound treatment, ML336 dissolved in DMSO was added to reaction mixtures along right before incubation at 37 °C at the indicated concentrations. The final DMSO concentration was 0.25%.

### Autoradiography of viral RNA

After extraction of viral RNA from the *in vitro* reactions, the RNA was mixed 1:1 with a glyoxal loading buffer/dye with ethidium bromide (Ambion) and denatured at 50 °C for 30 minutes. Samples were chilled briefly and loaded for RNA electrophoresis. RNA electrophoresis was performed through a denaturing agarose gel containing 0.8% agarose, 1X MOPS, and 2.2 M formaldehyde. Electrophoresis was performed at 60 V for 70 minutes, then the gels were rinsed in nuclease-free water once and treated with 0.1N NaOH for 40 minutes at room temperature with continuous rocking. Gels were neutralized in 20X saline-sodium citrate (SSC) buffer (3 M NaCl, 300 mM trisodium citrate, pH=7.0) for 40 minutes. RNA was transferred to a neutral nylon membrane. The RNA on the membrane was then UV cross-linked for 5 minutes at 4 mW/cm^2^. Autoradiograms were developed using a phosphor screen (Kodak K screen) and documented using a Pharos FX plus (BioRad).

### Cell-based anti-VEEV assay

We measured cell death using a CPE assay as previously described (9). Briefly, Vero 76 cells seeded in a 96 well plate were infected with VEEV TC-83 at an MOI of 0.05 in the presence of test compounds serially diluted to 8 different concentrations. Infected cells were incubated for 48 h and cell viability, protection from VEEV-induced CPE, was measured using CellTiter-Glo (Promega). EC_50_ was calculated with a 4-parameter logistic model.

### Statistics

Statistics were performed in GraphPad Prism version 7. Unless otherwise indicated significance was calculated using ANOVA with Dunnett corrections for multiple comparisons.

## Results

### TC-83 develops resistance to ML336 primarily through a mutation in nsP4

Previous work with our initial hit compound (PubChem CID: 15997213) has shown that VEEV strain TC-83 is able to develop resistance to compound treatment with mutations in nsP2 and nsP4 (10). To determine if ML336 treatment results in similar genomic changes conferring resistance, TC-83 was serially passaged in Vero76 cells in the presence of increasing concentrations of ML336. As shown in Figure 1 B, four biological replicates were analyzed (R1, R2, R3, R4) at each passage. Nucleotide changes that showed 10% frequency or less were generally not consistent between replicates. Only one nucleotide change led to a change in amino acid in TC-83 over passaging; Q210R/K in nsP4, which was fixed in all four biological replicates.

### VEEV viral RNA production peaks at 6-8 HPI

Resistance studies with our initial hit compound as well as with ML336 have shown that VEEV mutants can be selected for resistance to compound treatment. These mutations mapped within the N terminal domains of nsP2 and nsP4. These proteins are both involved in synthesis of viral RNA (13, 14). This lead to our hypothesis that ML336 inhibits VEEV replication by interfering with viral RNA synthesis.

VEEV has a very rapid replication cycle, with significant viral progeny being produced as early as four hours post-infection (HPI) in cell culture models (30). To determine the optimal time point to examine the antiviral activity of ML336 on RNA synthesis, first, we measured the rate of VEEV viral RNA synthesis and the amount of nsP2 over the course of infection using metabolic RNA labeling and western blotting. The synthesis of viral RNA was tracked in two-hour increments beginning at 2 HPI by pulsing cells with 3HU in the presence of act D, which allowed us to detect only viral RNA by inhibiting cellular RNA transcription from DNA template. We found that VEEV RNA synthesis was detectable at two HPI, and continued until 18 HPI (Fig 2 A). VEEV RNA synthesis reached its peak between 6 and 8 HPI and decreased at 10 HPI.

**Figure 2.**
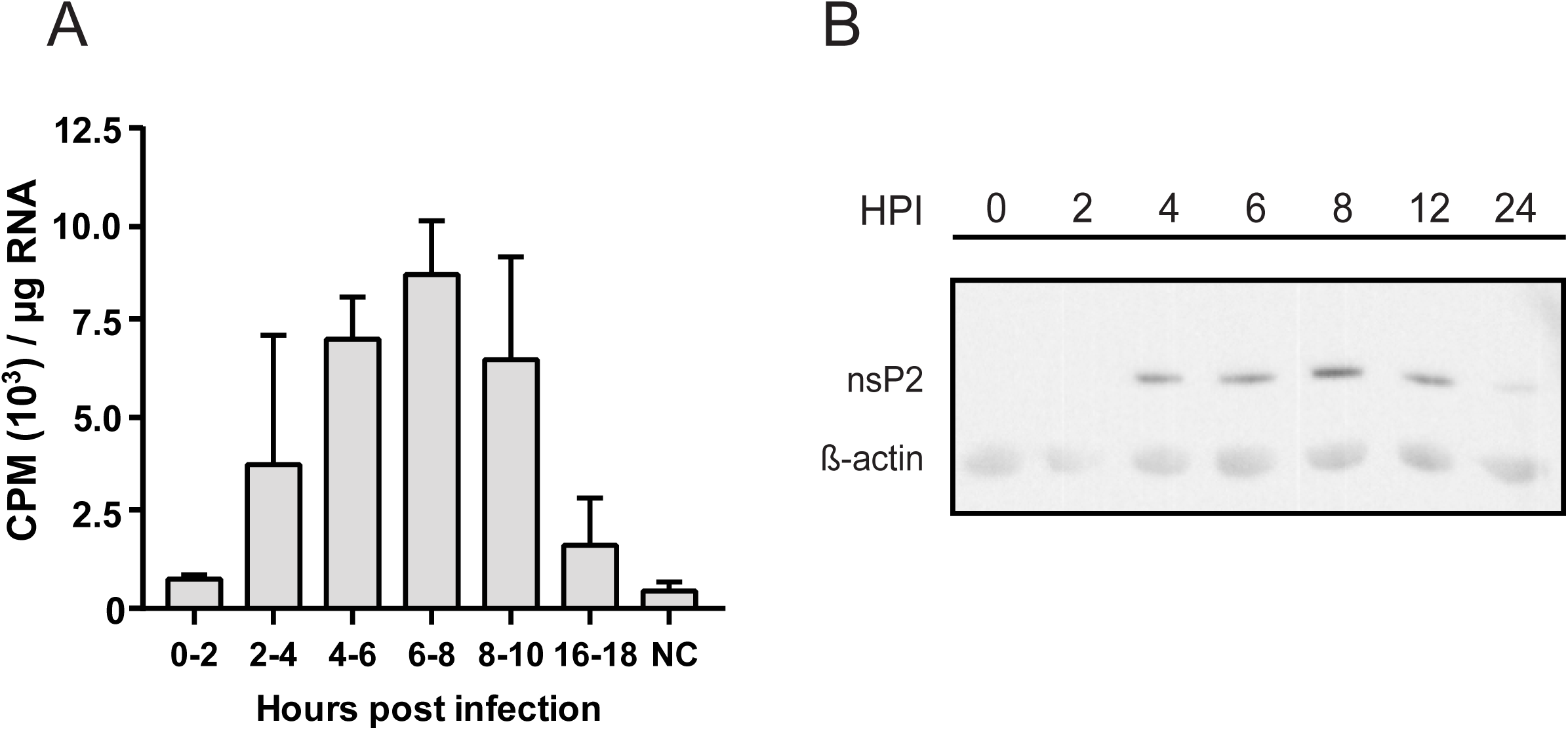
VEEV RNA production is maximal from 6 to 8 hours post infection. A) The amount of 3HU incorporated into viral RNA. Infected cells were incubated with 3HU and actD for two hours and the total RNA was isolated. The incorporated 3HU in the RNA was measured with a liquid scintillation counter. 3HU incorporation was normalized to µg of total RNA. VEEV viral RNA synthesis rate was maximal from six to eight HPI. B) Western blotting analysis probing for nsP2 in infected cells. Maximal nsP2 expression was achieved by 8 hours post infection.

The RNA synthesis data was compared to the amount of nsP2 determined by western blotting. We found that nsP2 was detectable beginning at 4 HPI and the levels peaked at 8 HPI. NsP2 was detected until 12 HPI with no change in the amount present (Fig 2 B). These data together indicate that the most active period of viral RNA synthesis of VEEV is from six to eight HPI. This time period correlates to the expression of nsP2 as measured by western blot, which is an important viral enzyme involved in the replication of VEEV RNA.

### ML336 inhibits viral RNA synthesis in infected cells

To test our hypothesis, the inhibitory effect of ML336 on viral RNA synthesis was measured in VEEV-infected cells using the 3HU-incoporation assay. 3HU was added to VEEV (strain TC-83)-infected cells in the presence of various concentrations of ML336 at 6 HPI, based on our data from the time-course experiment (Fig 3 A). The amount of 3HU incorporated into the total isolated RNA was measured and a dose response curve was generated to calculate the IC_50_ (Fig 3 A). ML336 showed strong, dose-dependent inhibition of viral RNA synthesis activity with an IC_50_ of 1.1 nM with a standard deviation (SD) of 0.7 nM and a 93% decrease in viral RNA synthesis at 40 nM. This data indicates that ML336 is a potent VEEV RNA synthesis inhibitor. The IC_50_ of ML336 in this viral RNA synthesis assay was comparable to the EC_50_ value of ML336 in the CPE-based assays (EC_50_ = 32 nM), indicating that RNA synthesis inhibition is likely to be a primary mechanism of ML336 action.

**Figure 3.**
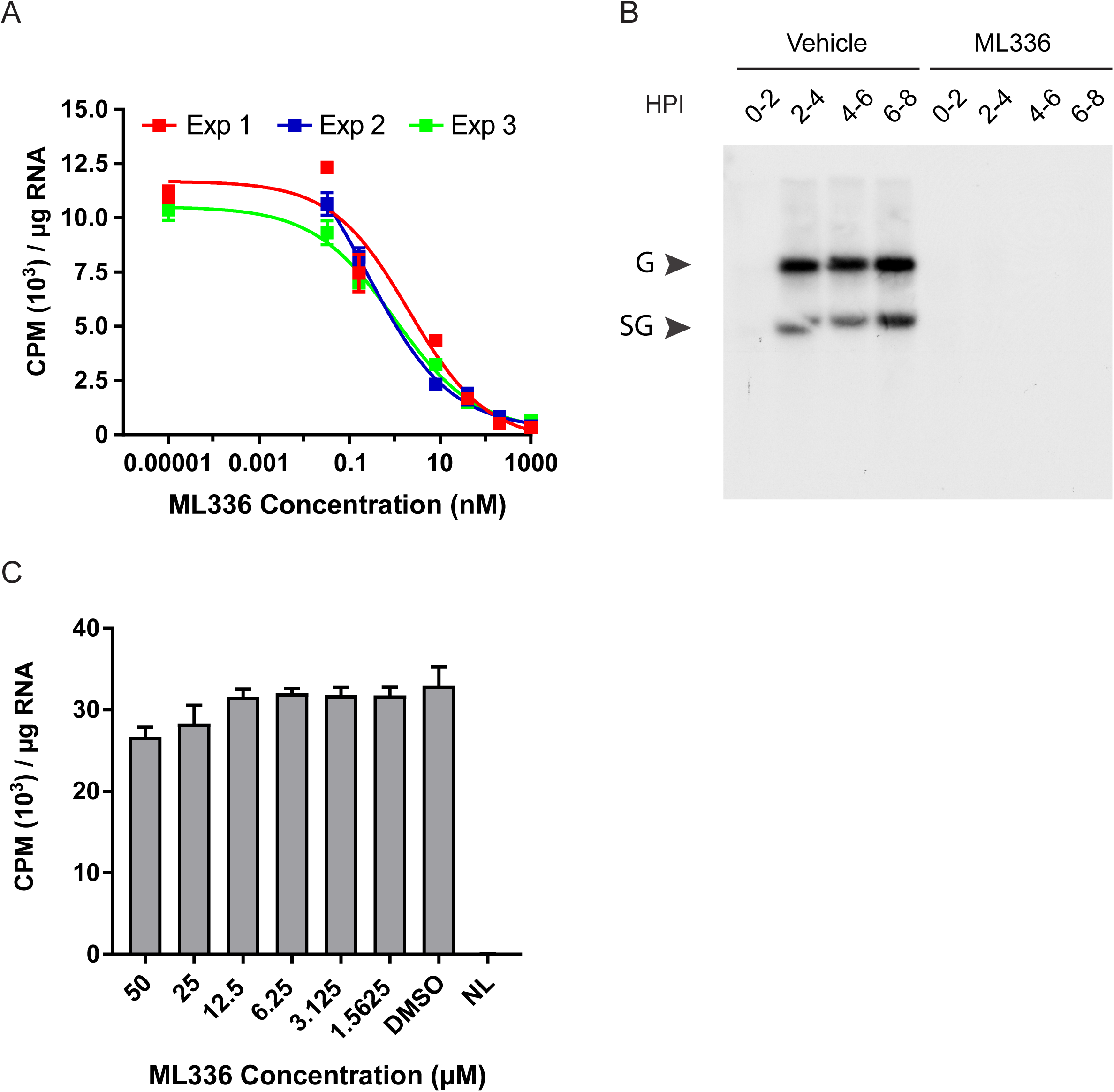
ML336 Inhibits the RNA synthesis of VEEV. A) A dose response curve was generated using 8 different concentrations of ML336 from 1 µM to 320 pM. 3HU incorporation was normalized to µg of total RNA. Non-linear regression was performed using in GraphPad Prism using a log vs response equation with a variable slope with four parameters. Lines were calculated for each individual experiment. The IC_50_ of each experiment was individually calculated and averaged for a final IC_50_ which is 1.1 nM with a standard deviation of 0.7 nM. B) Fluorgram of VEEV RNA with either ML336 treatment or DMSO control. Cells that were treated with ML336 show no production of either genomic or subgenomic viral RNA when compared to control. C) Uninfected BHK cells were incubated with ML336 at 6 concentrations and 3HU in the absence of act D. NL is negative control without 3HU.

To understand whether the inhibition of viral RNA synthesis was specific to genomic (49S) or subgenomic (26S) viral RNA, we visualized the viral RNAs that were produced in the presence/absence of ML336 using fluorography to detect 3HU-labeled viral RNA. As Figure 3 B shows, the addition of ML336 to infected cells at any time post infection up to 8 hours completely abrogated synthesis of both genomic and subgenomic viral RNA.

To confirm that RNA synthesis inhibition by ML336 is virus-specific and that there is no inhibition of cellular RNA synthesis, we investigated the effect of ML336 on cellular RNA synthesis. Uninfected BHK-21 cells were incubated with ML336 at the indicated concentrations or with a DMSO control in the presence of 3HU without act D (Fig 3 C). There was a small decrease in cellular RNA synthesis only at the highest dilutions, 50 µM and 25 μM, indicating that the inhibitory activity of ML336 against cellular RNA synthesis is negligible.

These data clearly indicate that ML336 strongly inhibits viral RNA synthesis. ML336 was effective against both the 49S genomic and 26S subgenomic RNAs. The effect of ML336 is specific for viral RNA and had only minimal effects on the synthesis of cellular RNA. These findings suggest that the primary mechanism of action of ML336 is viral RNA synthesis inhibition.

### ML336 inhibits the activity of the mature replicase complex

The experiments above demonstrated that ML336 inhibits VEEV viral RNA synthesis; however, it was not clear which stage of RNA synthesis (positive sense and/or negative sense RNA synthesis) was inhibited by ML336. For alphaviruses, the synthesis of positive strand RNA is tightly regulated by negative RNA synthesis. The positive sense RNA, which is produced by the mature replicase complex, nsP1/2/3/4 (14), is dependent on the presence of negative sense genomic RNA which is the template for positive strand synthesis. Therefore, the inhibition of negative sense RNA synthesis would result in loss of positive sense RNA synthesis. To determine if the inhibition of viral RNA synthesis by ML336 requires the inhibition of negative sense RNA synthesis, we analyzed the RNA synthesis inhibitory activity of ML336 in the presence or absence of cycloheximide in our radiolabeling assay. The negative sense RNA is produced only by the newly translated polyprotein, nsP123/4 (22, 23). CHX inhibits the production of new protein which will prevent the production of nsP123/4 and thus negative strand RNA (14, 31). So, the addition of CHX to our RNA synthesis assay allows us to measure positive sense RNA synthesis by the pre-formed viral replicase complex.

Figure 4 A shows amount of 3HU labeled RNA that was produced in the presence and absence of CHX and ML336. Treatment with CHX showed no significant difference from the positive control, indicating there is no significant production of negative sense RNA at this time. Treatment with ML336 with or without CHX equally abrogated viral RNA production. This result showed that at this stage in infection ML336 inhibited the synthesis of positive sense RNA, which is generated by the mature replicase complex. This data supported our hypothesis that ML336 inhibits the RNA synthesis activity of the mature, fully formed replicase complex.

**Figure 4.**
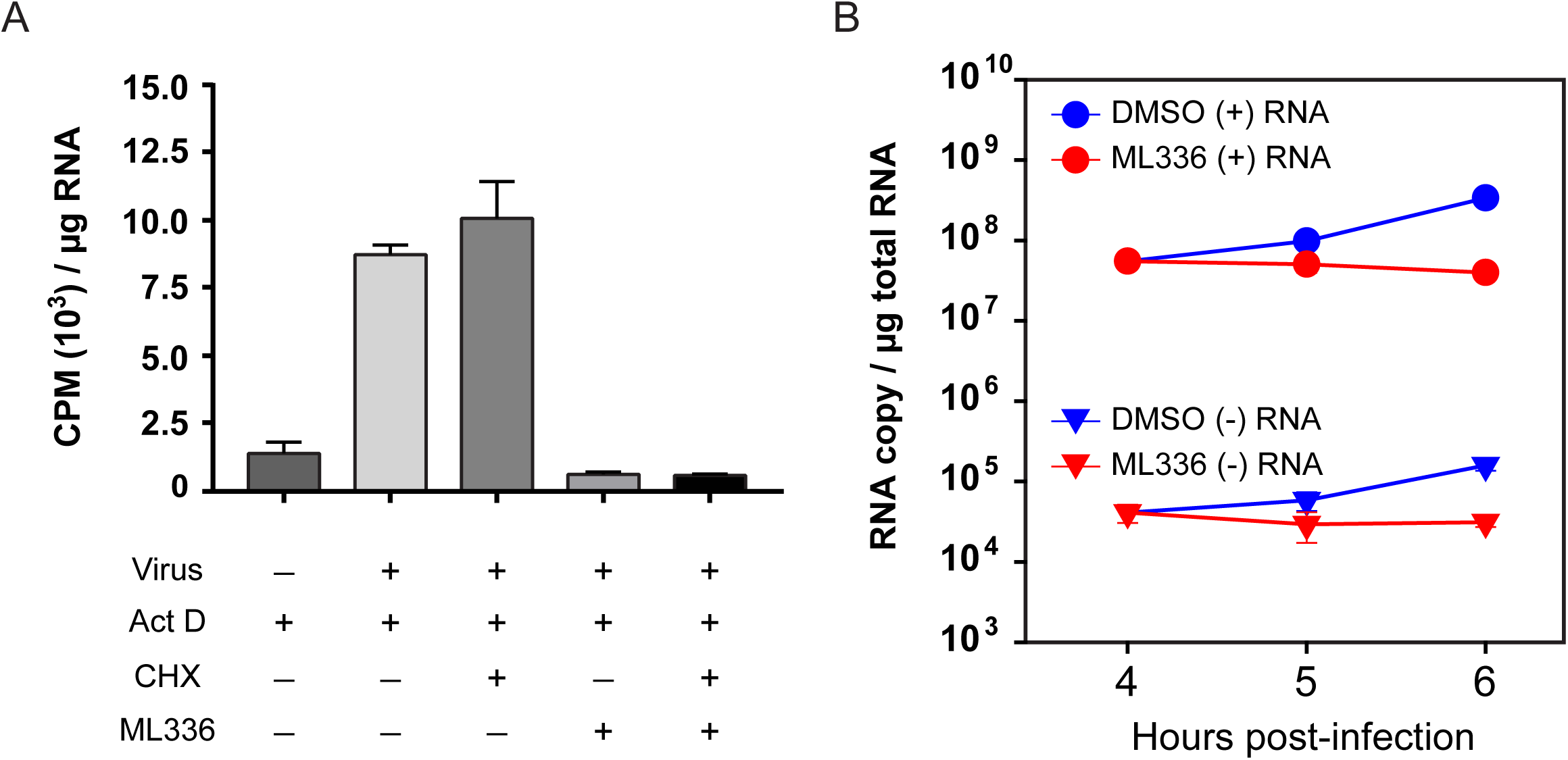
ML336 inhibits all stages of VEEV RNA production. A) Infected cells were treated with different combinations of CHX, act D, and ML336 as indicated. RNA was isolated and 3HU incorporation was quantified as described above. B) Strand-specific qRT-PCR was carried out on RNA extracted from VEEV infected cells at the time points indicated. Infections were carried out in the presence of either ML336 or DMSO vehicle control. The presence of ML336 during infection prevented the production of both positive and negative strand RNA.

### ML336 is a potent inhibitor of both positive and negative RNA synthesis of VEEV

Our data has shown that ML336 interferes with the production of VEEV RNA, and ML336 efficiently inhibits the activity of the mature replicase complex, which is responsible for the production of VEEV positive sense RNA. While there is much less negative sense viral RNA present during infection, the negative strand is critical for RNA replication because it is the template that is used to create more positive sense RNA (14). To determine if ML336 was active against negative strand synthesis as well, we used a strand-specific qRT-PCR (28). ML336 or DMSO were added to VEEV-infected cells at 4 HPI, and the positive and negative viral RNA copy numbers were quantified and compared. As shown in Figure 4 B, the DMSO control group had almost a 100-fold increase in the amount of positive-sense RNA, while there was approximately a 50-fold increase of negative-sense RNA.

In the presence of ML336, there was no change in the amount of positive-sense viral RNA from 4-6 HPI. Similarly, when ML336 was added there was no increase in the amount of negative sense RNA. The presence of ML336 did not affect the ratio of positive sense to negative sense RNA. Positive sense RNA was detected at levels 10,000-fold higher than the negative sense RNA at all time points and in both ML336 treated and control reactions. This difference in the levels of the RNA species has been reported previously in alphaviruses (32). These results show that ML336 inhibits the synthesis of both the positive and negative sense strands of VEEV RNA.

### RNA synthesis inhibition is the primary mechanism of action of ML336

To further validate that inhibition of viral RNA synthesis is the mechanism of action of ML336, we employed analogues of ML336 in the 3HU incorporation assay. We hypothesized that if inhibition of viral RNA synthesis was the primary MOA of the antiviral effect of this class of compounds, then their viral RNA synthesis inhibitory activity would correlate with antiviral potency.

Nine compounds with 50% CPE inhibitory concentration (EC_50-CPE_) ranging between 0.1 μM and > 50 μM were tested at 1 µM in the 3HU labeling assay and their inhibitory activities were compared with their EC_50-CPE._ We also tested an antiviral compound, ML416, of which the main antiviral mechanism is unrelated to viral RNA synthesis inhibition (Fig 5 A) (8). In this analysis, while ML416 did not show significant viral RNA synthesis inhibition, all of the analogue compounds similar to the ML336 showed significant inhibition of viral RNA synthesis. Moreover, we found a relationship between the antiviral activity of a compound and the degree of RNA inhibition in the 3HU incorporation assay. This result strongly suggested that RNA synthesis inhibition is a major mechanism of antiviral action for compounds based around this amidine scaffold.

**Figure 5.**
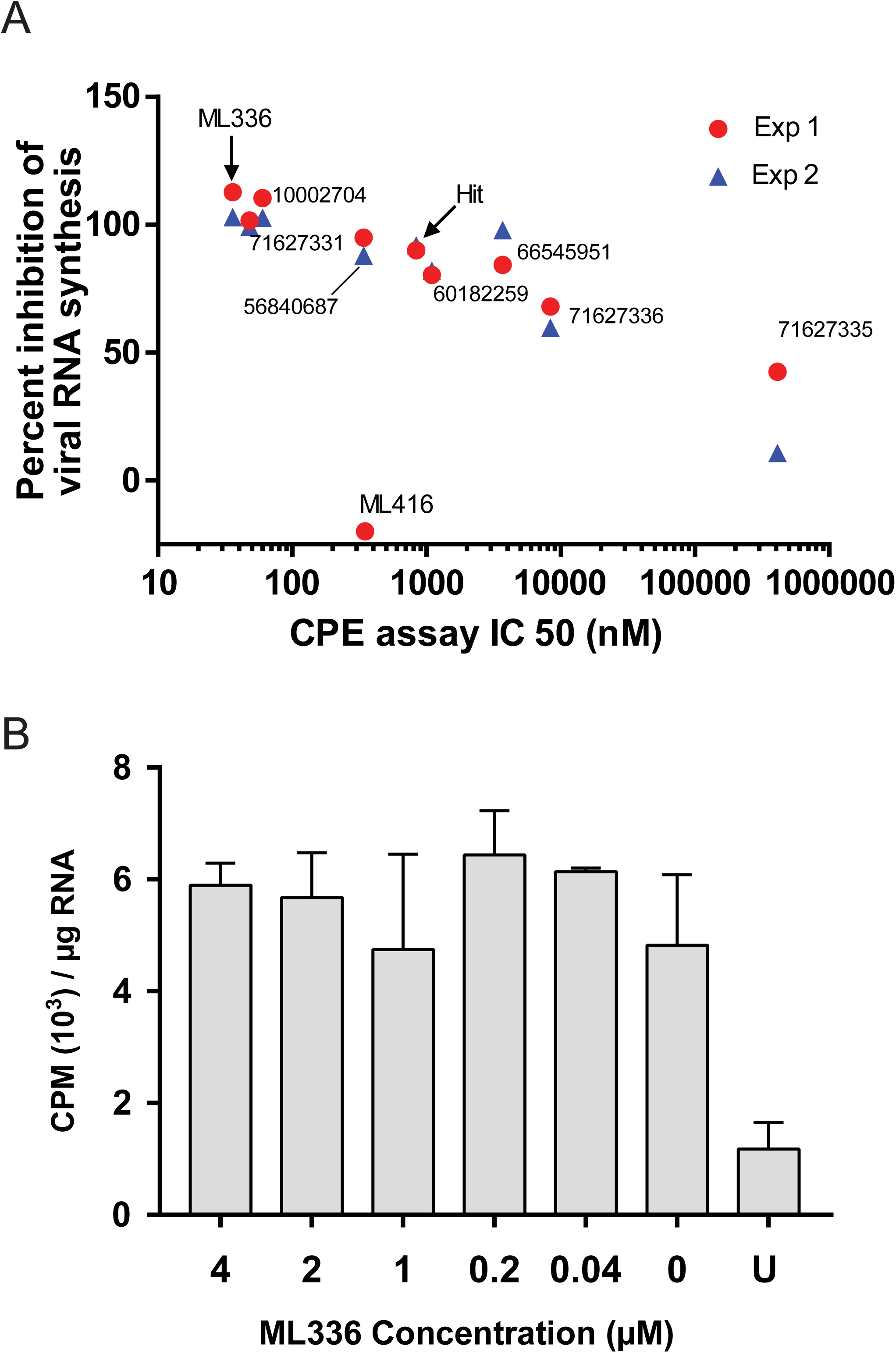
Inhibition of RNA synthesis is a primary mechanism of action of ML336. A) Anti-VEEV potency of ML336 analogue compounds was compared with their anti-viral RNA synthesis efficacy. Compounds highly effective at inhibiting RNA synthesis showed lower IC_50_ values in the CPE assay. ML416, which is an antiviral with no effect on viral RNA synthesis, was included in the assay as a control. B) The anti-viral RNA synthesis effect of ML336 was specific to VEEV, no effect to CHIKV RNA synthesis. CHIKV RNA synthesis was measured in the same 3HU incorporation assay that had been used for VEEV in the presence of six different concentrations of ML336 as indicated. There was no effect on CHIKV at any of the concentrations examined. NC is an uninfected negative control.

### ML336 has no effect on CHIKV RNA synthesis, consistent with its lack of anti-CHIKV activity

Previously we determined that compounds based around the ML336 scaffold had no antiviral effect on CHIKV in cell culture assays (33). Therefore, we hypothesized that ML336 would not show any inhibitory effect on CHIKV RNA synthesis if viral RNA synthesis inhibition was the major antiviral MOA of the compound against VEEV. To test the effect of ML336 on CHIKV RNA synthesis, we used the same 3HU incorporation assay format as previously described with CHIKV in place of VEEV. As can be seen in Figure 5 B, ML336 did not show any significant effect on CHIKV RNA synthesis even at 4 µM, *P* > 0.22, ANOVA, the highest concentration we tested and nearly 1000-fold higher than the IC_50_ value of the compound against VEEV in this assay, compared to an untreated positive control.

These results indicated that there is no effect of ML336 treatment on CHIKV RNA synthesis. This finding is consistent with previous results showing that ML336 has no antiviral effect against CHIKV in a cell culture model. This result supports our conclusion that RNA synthesis inhibition is the major mechanism of the antiviral activity of ML336.

### ML336 inhibits VEEV RNA synthesis in a cell-free system

Our genetic and biochemical studies have indicated that ML336 was likely to directly interact with the viral replicase complex to inhibit viral RNA synthesis. To determine if ML336 directly interacted with the replicase complex, we employed a cell-free viral RNA synthesis assay (29). This cell-free system uses the membranous fraction of infected cells (the P15) which is enriched for viral replicase complexes. This method has been validated and used for multiple alphaviruses (34). The use of the P15 provides both template and polymerase in an *in vitro* reaction to produce viral RNA (18).

As shown in Figure 6, while there is no radiolabeled RNA produced by the P15 fraction of uninfected cells, three distinct viral RNA bands corresponding to 26S, 49S, and a replication intermediate were present in the positive control. The addition of ML336 in the reaction decreased the amount of radiolabeled RNA. This indicated that ML336 is inhibiting the synthesis of viral RNA in this cell-free assay, supporting our hypothesis that ML336 directly inhibits viral RNA synthesis.

**Figure 6.**
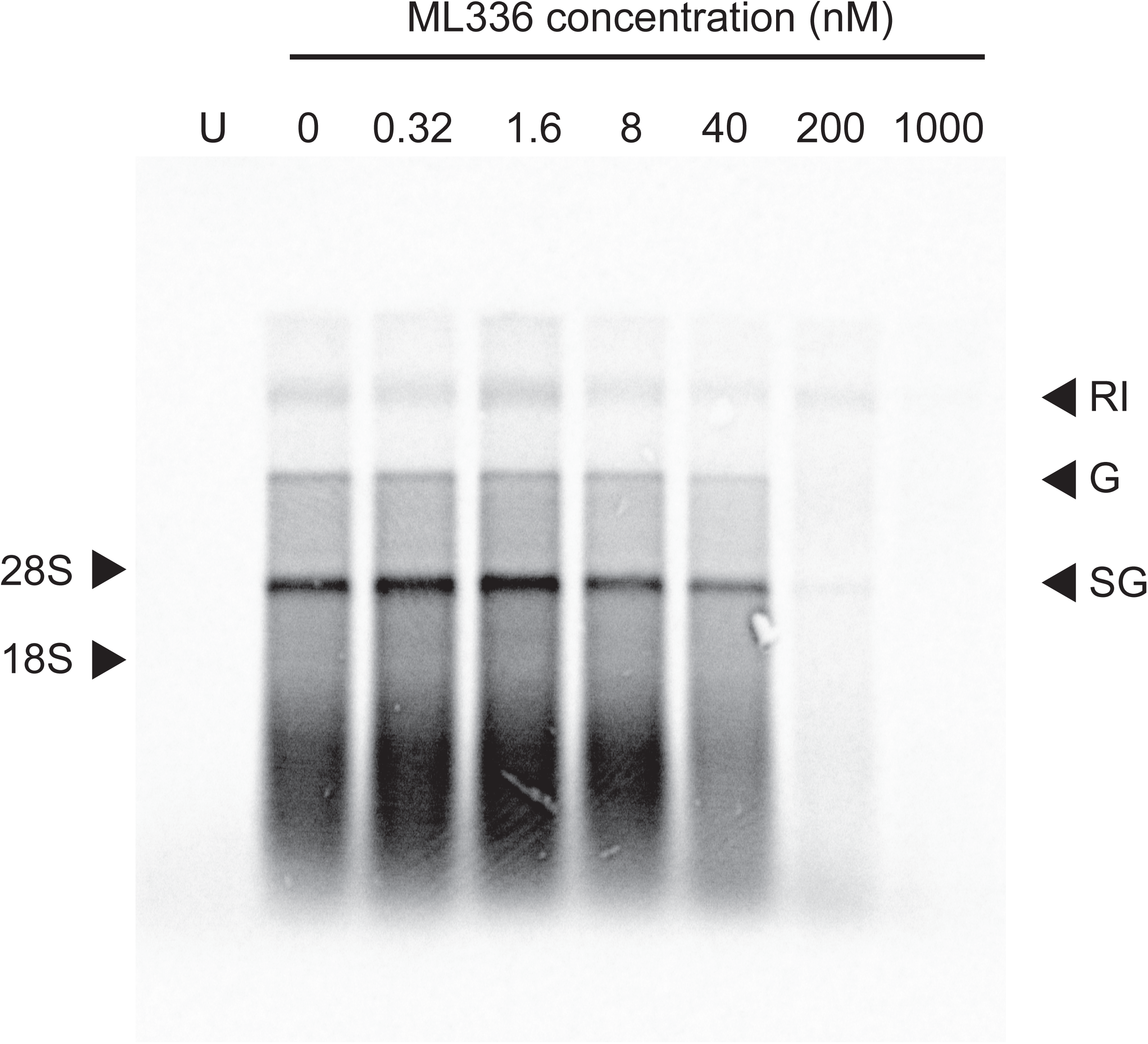
ML336 inhibits VEEV RNA production in a cell-free system. P15 fractions isolated from either infected or uninfected (U) control cells were incubated with [α-^33^P]-CTP in a reaction buffer described in Material and Method. As the concentration of ML336 increased, the amount of [α-^33^P] labeled viral RNA decreased for all RNA sizes. Replication intermediate, genomic, and subgenomic RNA are denoted as RI, G, and RG, respectively.

Taken together, these data indicate that ML336 inhibits VEEV by directly preventing viral RNA synthesis. ML336 was efficacious in both cell-based and cell-free viral RNA synthesis assays, and the compound showed strong inhibitory activity against both positive and negative-sense RNA synthesis. Our data support the conclusion that the novel compound, ML336, inhibits VEEV replication by interfering with viral RNA synthesis

## Discussion

VEEV is an alphavirus that circulates in South and Central America cycling between mosquito vectors and small mammal hosts (3). VEEV spontaneously and sporadically has caused large outbreaks of both human and equid infection, leading to over 109,000 human infections with over 500 associated fatalities in the decade from 1962 to 1972 (3). This virus is also regarded as a Select Agent by the Centers for Disease Control and Prevention due to the high risk of VEEV being used as an agent of bioterrorism or biological warfare when it is aerosolized (4, 35). Despite the ongoing risk VEEV poses to public health, there are currently no approved therapeutics or preventatives for VEEV infection in humans. This makes development of potential VEEV inhibitors a top priority.

Our lab has previously published on both our discovery of an anti-VEEV hit compound (CID: 15997213), and the development of our current lead amidine compound, ML336 (9-11). However, until now, the MOA of these compounds was unknown. Our initial attempts using resistant viruses identified mutations in the N terminal regions of both nsP2 and nsP4. These regions currently have no annotated function; however, there are near areas of these proteins that are necessary for viral RNA synthesis. The mutations in nsP2 are proximal to a known helicase domain, and those in nsP4 are near to the RNA dependent RNA polymerase domain (19, 20). We have confirmed that resistance mutations also arise in nsP4 in response to ML336 treatment. This information led to our hypothesis that ML336 inhibits viral replication by interfering with VEEV RNA synthesis.

To determine if viral RNA synthesis is inhibited by ML336, we utilized a radiolabeling assay using 3HU. This assay allowed us to detect only the production of new viral RNA separately from previously synthesized RNA in the infected cells. Using this assay, we found that ML336 indeed inhibits the production of VEEV RNA. This RNA inhibition was also highly efficient (EC_50_ = 1.1 nM), even more so than the initial CPE inhibition described in our initial cell-based assay (Fig 3).

This RNA inhibition effect was also present in ML336 analogs that showed potency against VEEV CPE, indicating a relationship between anti-CPE activity and RNA synthesis inhibition. We had also seen previously that ML336 has no effect on a representative Old World alphavirus, CHIKV, in our CPE-based assay. We confirmed that ML336 has no effect on the RNA synthesis activity of CHIKV, even at 4 µM, a concentration 4,000-fold higher than the IC_50_ value of ML336 against VEEV in this assay. These data showed that 1) the ML336 scaffold possesses viral RNA synthesis inhibition activity, and 2) the RNA synthesis inhibition activity of the ML336 scaffold correlated well with antiviral activity in cells, strongly supporting the conclusion that the major antiviral mechanism of action of these compounds is viral RNA synthesis inhibition. Our argument is also supported by the fact that ML336 did not inhibit the RNA synthesis of CHIKV, a virus for which ML336 has no antiviral effect.

In this paper, we show that the inhibition of RNA synthesis by ML336 extends to all viral RNA species that are produced during replication. Using fluorography, we showed that ML336 inhibited the synthesis of both the 49S genomic and the 26S subgenomic RNA strands. Further, by employing CHX, which inhibits the translation of new nsP polyprotein and thus inhibits the synthesis of the negative-sense RNA, we demonstrated that ML336 directly inhibited the activity of the pre-formed, mature replicase complex, and did not require the inhibition of negative sense RNA synthesis. Finally, using strand specific qRT-PCR we were able to show that ML336 prevented the synthesis of both positive and negative sense RNA. All of these data collectively indicate that ML336 effectively inhibits the synthesis of every class of RNA that is synthesized by VEEV over the course of infection.

Based on the MOA, antiviral inhibitors can be classified into two categories; 1) direct-acting antivirals which act directly on viral components, and 2) host-targeting antivirals which rely on agonizing or antagonizing cellular processes. While host-targeting antivirals, such as ribavirin and interferon, are often effectively used in the clinic for many viral diseases, they also often have severe side effects due to off target interactions in the cell (36-39). Due to these significant side effects, much current research is focused on the development of direct acting antiviral molecules, in the hope that they will have fewer side effects. Importantly, VEEV is not sensitive to the treatment with ribavirin (6).

To determine if ML336 was acting directly on viral proteins, it was necessary to employ an assay to evaluate inhibitory activity against viral RNA production in a cell-free environment. For alphaviruses, this poses some challenges, due to the difficultly of ectopic expression the nsPs. The nsPs, as the functional whole replicase complex, are known to be difficult to express in common vector systems. NsP4 is also difficult to express in isolation (20). Currently no three-dimensional structures of the replicase complex are available (40). To evaluate the anti-RNA synthesis activity of ML336 in a cell-free, biochemical assay system, we used the P15 of infected cells, as has been reported previously with Old World alphaviruses (13). The P15, the intracellular membranous fraction, from alphavirus-infected cells is known to contain most of the replicative activity (25, 29). Using this method, we were able to develop a cell-free assay to investigate the activity of the VEEV viral replicase complex. The P15 fraction displayed similar RNA synthesis activity as Old World alphaviruses, with a slight difference in the optimal Mg^++^ concentration of 12 mM (data not shown), instead of 3 mM as described in Albulescu’s paper (34). In our assay, 49S genomic, 26S subgenomic, and replication intermediates were detected.

The RNA synthesis activity of the P15 fractions were significantly inhibited when incubated in the presence of ML336. The synthesis of all species of viral RNA was equally affected as detected by autoradiography. The potent inhibition of viral RNA synthesis by ML336 in a cell-free assay indicates that the compound does not likely target a cellular protein, as a majority of the cellular components had been removed. This leads to our conclusion that ML336 is acting directly on the viral replicase complex to specifically limit the production of viral RNA.

Our data clearly showed that ML336 is able to inhibit the synthesis of VEEV RNA, and that this inhibition is a primary mechanism of the antiviral activity of this compound. However, it is still unknown which of the activities of the non-structural proteins are being affected by ML336. ML336 inhibits both negative sense RNA synthesis, carried out by newly synthesized nsPs, and positive sense RNA synthesis, which is carried out by the mature replicase complex (Fig 4). The viral replication machinery has a presumably different structure during these stages, due to the cleavages that occur in the polyprotein before the formation of the replicase complex (21). This indicates that ML336 is likely acting in a manner that is independent from the spatial relationship of the nsPs. If the activity of ML336 was dependent on interfering with the formation of the nsP complex, this RNA synthesis inhibition activity would have no effect on the function of pre-formed replicase complexes. However, data from our experiment using CHX (Fig 4) clearly showed that ML336 does inhibit the activity of pre-formed replicase complexes.

This inhibition of both newly synthesized polyproteins, and mature replicase complexes leads to our hypothesis that ML336 is able to disrupt the molecular activities of nsP4, and is able to inhibit RNA elongation. Nsp4 is the active RNA dependent RNA polymerase of VEEV, responsible for both positive and negative sense RNA synthesis (20). Thus, if ML336 were to inhibit the activity of nsP4, all RNA replication would stop, regardless of the stage of infection.

This hypothesized interaction with nsP4 is also supported by the occurrence of mutations in this protein that confer resistance to ML336. To establish the direct antiviral action of ML336, a direct interaction between ML336 and the replicase complex should be investigated. This is of particular importance to confirm the residues where this interaction occurs. Understanding of such an interaction could also allow us to use ML336 as a biological probe to understand currently unknown biological functions of the nsPs. In addition, detailed understanding of the interaction between the compound and the replicase complex would allow us to develop the ML336 scaffold for direct antiviral therapeutics for Old World alphaviruses such as CHIKV as well.

We have also investigated the use of steady state enzyme kinetics as a way to determine the general action of ML336; if it is a competitive, noncompetitive or mixed inhibitor of the RNA synthesis machinery. However, these experiments are currently not feasible, primarily due to the complexity of our cell-free mixture, which is an isolation of an entire cellular compartment.

Collectively our data show that ML336 is a small molecule that inhibits VEEV replication by interfering in the production of viral VEEV RNA. Viral RNA synthesis is reduced regardless of polarity, and both genomic and subgenomic RNA synthesis are affected. We found a high degree of correlation between RNA synthesis inhibition activity and viral replication inhibition with analogue compounds around the ML336 scaffold, strongly suggesting that the inhibition of RNA synthesis is the major mechanism of the antiviral activities of these compounds. Inversely, ML336 did not show any RNA synthesis inhibitory activity for CHIKV, which resisted the treatment of ML336 in the antiviral assay. Most excitingly, ML336 displayed strong anti-viral RNA synthesis activity even in a cell-free system, indicating a direct interaction between the compound and the viral replicase complex. Our results show that ML336 has exciting potential as a lead scaffold compound for a new class of anti-alphaviral drugs and could be used as a molecular probe in further characterization of the alphaviral nsPs.

